# Functional Data Analysis of Spatial Clustering Identifies Prognostic T Cell Patterns in Ovarian Cancer

**DOI:** 10.64898/2026.07.02.735980

**Authors:** Chase J. Sakitis, Daisy Liao, Brett M. Reid, Mary K. Townsend, Joellen M. Schildkraut, Andrew B. Lawson, Shelley S. Tworoger, Kathryn L. Terry, Lauren C. Peres, Julia Wrobel, Alex C. Soupir, Brooke L. Fridley

## Abstract

Spatial proteomic imaging technologies enable the simultaneous assessment of immune cell abundance and spatial organization within the tumor microenvironment. Spatial clustering is commonly summarized using measures such as Ripley’s *K* or nearest`Ö`neighbor *G*-functions at a fixed radius. However, these approaches depend on scale selection and may obscure biologically relevant patterns occurring across spatial ranges. We propose a functional data analysis (FDA) framework to model spatial clustering trajectories derived across a continuum of radii. Functional principal component analysis (FPCA) was used to summarize dominant modes of spatial variation, and resulting scores were incorporated into Cox proportional hazards models as both main effects and interaction with immune cell abundance. The approach was applied to multiplex immunofluorescence data from five ovarian cancer studies, comprising 773 high`Ö`grade ovarian serous tumors. Analyses focused on CD3+ and CD8+ T cell populations within the tumor compartment of the tissue, adjusting for age at diagnosis and cancer stage, with study-specific estimates combined using random-effects meta-analysis. Higher abundance of both T cells and CD8+ T cells was consistently associated with improved overall survival. Beyond abundance, spatial features captured by the leading functional principal component were independently associated with survival, particularly for CD8+ T cells. Interaction models further showed that the prognostic effect of immune infiltration depended on spatial clustering, with tumors characterized by high abundance and low spatial clustering exhibiting the most favorable outcomes. These findings indicate that spatial organization provides complementary prognostic information beyond abundance alone and suggests that more diffuse immune infiltration may reflect more effective anti-tumor activity in ovarian cancer. Overall, FDA offers a flexible and interpretable framework for modeling spatial clustering across scales and identifying prognostic spatial features not captured by fixed-radius or distance analyses.

## 1. Introduction

Characterizing the tumor microenvironment (TME) is essential for cancer research and improving therapeutic outcomes^1–3^. A detailed understanding of the cellular, molecular, and spatial interactions within the TME can reveal mechanisms of tumor progression, immune evasion, and treatment resistance^1,4–6^. Spatial proteomic imaging data has transformed how the TME is studied in cancer research^7,8^. Not only can researchers assess differences in cell population abundances between conditions or clinical outcomes^8^, but they are also able to study the spatial architecture of different cell populations^7^.

Epithelial ovarian cancer (EOC) presents a highly heterogeneous and immunologically active TME that plays a critical role in disease progression and patient outcomes^9,10^. Tumor-infiltrating lymphocytes (TILs) are frequently observed within tumor islets and have been consistently associated with improved survival, either alone^10–12^ or in conjunction with B-cells^13^. However, the immune landscape is complex, as immunosuppressive populations such as regulatory T cells and tumor-associated macrophages contribute to immune evasion and poorer clinical outcomes^14,15^. Despite the active TME in ovarian cancer, the efficacy of immunotherapy has been modest compared to other malignancies^16^ and limited to a small subset of patients^17^. However, recent clinical studies combining pembrolizumab and paclitaxel demonstrated improved outcomes in ovarian cancer^18,19^, with emerging evidence suggesting that spatial features of the TME, such as reduced clustering of CD8+ T cells among responders, may provide additional insight beyond immune cell abundance alone. Collectively, these findings underscore the importance of spatial context in shaping both prognosis and therapeutic response in EOC.

Prior studies of spatial clustering or co-localization of different cell populations in the TME have often used Ripley’s *K*^20^ or nearest-neighbor *G* function^21^ with a set radius for their association analysis^22–24^. However, these measures are computed at various radii to assess the clustering at different spatial ranges^22,23,25^. This can be problematic, as identifying the spatial radius that is most biologically relevant is inherently uncertain. Instead of selecting a pre-specified radius for the association of the level of spatial clustering with clinical outcomes, we propose the use of functional data analysis (FDA) and analysis of the spatial clustering curves or spatial trajectories to better understand the relationship between spatial clustering of immune cell populations in the TME and clinical outcomes. Spatial relationships between T cells (i.e., CD3+) and CD8+ T cells in the ovarian TME and survival was found in prior studies based on a fixed radius^26,27^. Here, our goals are to assess the utility of FDA for spatial analysis of multiplex immunofluorescence (mIF) data and replicated prior findings based on fixed radii analysis for the association between spatial clustering of T cells and ovarian cancer survival.

## 2. Methods

### 2.1. Ovarian Cancer Studies

#### African American Cancer Epidemiology Study (AACES)

AACES is a population-based case-control study of self-identified African American or Black women diagnosed with epithelial ovarian cancer across 11 U.S. geographic regions, with 592 cases and 752 controls enrolled between 2010 and 2015^28^. Detailed epidemiologic and clinical data were collected through structured computer-assisted telephone interviews, including demographic, reproductive, lifestyle, and medical history information. Formalin-fixed, paraffin-embedded (FFPE) tumor blocks were obtained, and tissue microarrays (TMAs) were generated by sampling multiple cores from each tumor. All tumors underwent centralized pathology review at Duke University to verify diagnosis and histologic subtype. Vital status and follow-up information were obtained through annual follow-up surveys and medical record abstraction, with ongoing follow-up initiated after enrollment. The study protocol was approved by the institutional review boards at Duke University Medical Center and participating study sites, including collaborating institutions across Alabama, Georgia, Louisiana, North Carolina, South Carolina, Tennessee, Texas, Michigan, Ohio, and New Jersey, as well as relevant state cancer registries and SEER registries^28^.

#### North Carolina Ovarian Cancer Study (NCOCS)

NCOCS is a population-based case-control study conducted between 1999 and 2008 across 48 counties in North Carolina, with 1,173 enrolled cases^29^. Baseline clinical and epidemiologic data were collected through in-person interviews administered by trained nurses. FFPE tumor blocks were obtained from the facility where debulking surgery was performed. Tumor tissue was available for 95% of cases, and all specimens underwent centralized pathology review to confirm diagnosis. Representative tumor regions were selected and incorporated into TMAs. Vital status was ascertained through the Social Security Death Index, the North Carolina Central Cancer Registry, and the National Death Index, with follow-up completed through 2020. The study protocol was approved by the institutional review boards at Duke University Medical Center and participating institutions involved in NCOCS.

#### New England Case Control Study of Ovarian Cancer (NECC)

NECC is a population-based case-control study conducted in Eastern Massachusetts and New Hampshire^30^. Incident epithelial ovarian cancer cases were identified between 1984 and 2008, yielding 3,083 eligible cases, of whom 2,203 (71%) participated. Detailed epidemiologic data, including reproductive, lifestyle, and clinical factors, were collected through in-person interviews. FFPE tumor blocks containing representative tumor material were obtained, and a gynecologic pathologist reviewed each specimen to confirm histology and tumor grade. Representative tumor regions were subsequently selected for TMA construction. Vital status was ascertained through linkage with cancer registries and the National Death Index. The study protocol was approved by the institutional review boards at participating institutions, including the Dana-Farber Harvard Cancer Center and Dartmouth Medical Center.

#### Nurses’ Health Study I (NHS) and II (NHSII)

NHS was initiated in 1976 and enrolled 121,700 female registered nurses aged 30–55 years^31,32^, while NHSII began in 1989 and enrolled 116,429 female registered nurses aged 25–42 years^33^. Demographic, lifestyle, and clinical data were collected prospectively through biennial questionnaires. For participants with confirmed ovarian cancer and available pathology reports, FFPE tumor tissue blocks were obtained. A gynecologic pathologist reviewed all specimens and selected tumor regions for inclusion in TMAs^34^. Vital status was ascertained through linkage with the National Death Index, state cancer registries, and ongoing cohort follow-up procedures^32,35^. The NHS and NHSII study protocols were approved by the institutional review boards of Brigham and Women’s Hospital, the Harvard T.H. Chan School of Public Health, and participating cancer registries, as required. Additional cohort details have been described previously^35^.

All analyses were restricted to samples from subjects with tumors classified as high-grade serous ovarian carcinoma (HGSOC). A summary of these participant characteristics for each EOC study is shown in **Table 1**. For all studies, FFPE ovarian tumor tissue samples were collected and used to create TMAs or select regions of interest (ROIs) from whole-slide tissue. mIF imaging was performed using the AKOYA Biosciences OPALTM 7-Color Automation IHC Kit. Following staining, image collection was performed by Vectra®3 Automated Quantitative Pathology Imaging System (0.499µm/pixel). InForm was used for spectral unmixing, and HALO was used for cell phenotype assignments. Generation of spatial mIF data for all studies was completed at the Advanced Analytical and Digital Laboratory under the Pathology Department at the Moffitt Cancer Center.

**Table 1:** Summary of the subjects and corresponding tumor samples included in the study.

|  |  |  |  |  |  |
| --- | --- | --- | --- | --- | --- |
|  | **AACES** | **NCOCS** | **NECC** | **NHS** | **NHSII** |
| **Number of Subjects** | 155 | 136 | 175 | 239 | 68 |
| **mIF Sample Type** |  |  |  |  |  |
| TMA – N Subjects | 62 | 0 | 175 | 239 | 68 |
| ROI – N Subjects | 93 | 136 | 0 | 0 | 0 |
| **Age at Diagnosis (SD)** | 59.9 (9.52) | 58 (8.94) | 59.1 (9.42) | 66.6 (9.64) | 53 (6.96) |
| **Stage (%)** |  |  |  |  |  |
| Early (Stage 1 or 2) | 39 (25.2%) | 23 (16.9%) | 33 (18.9%) | 43 (18%) | 22 (32.4%) |
| Late (Stage 3 or 4) | 116 (74.8%) | 113 (83.1%) | 142 (81.1%) | 196 (82%) | 46 (67.6%) |
| **Vital Status** |  |  |  |  |  |
| Alive | 54 | 24 | 44 | 31 | 38 |
| Deceased | 101 | 112 | 131 | 208 | 30 |
| **Follow-up Time After Diagnosis (in months)** |  |  |  |  |  |
| Median | 53.19 | 52.0 | 56.6 | 49.0 | 89.0 |
| 10<sup>th</sup>-90<sup>th</sup> percentile | 17.3 – 92.0 | 18.2 – 199.2 | 21.4 – 151.6 | 8.0 – 190.8 | 33.4 – 212.3 |
| **Number of Subjects with ≥6 positive cells** |  |  |  |  |  |
| CD3+ | 132 (85.2%) | 118 (86.8%) | 156 (89.1%) | 192 (80.3%) | 56 (82.4%) |
| CD3+CD8+ | 109 (70.3%) | 93 (68.4%) | 119 (68%) | 128 (53.6%) | 45 (66.2%) |

### 2.2. Spatial Summary Measures and Functional Data Analysis

Recent advances in multiplex spatial imaging technologies enable high-dimensional, single-cell characterization of protein expression while preserving spatial organization within intact tissues^36^. These platforms generate spatially resolved datasets consisting of individual cells with defined locations and phenotypes, providing the foundational inputs for downstream spatial modeling. Spatial analyses require a set of spatially resolved cells, the two-dimensional locations of those cells within each tissue sample, and discrete cell phenotypes defining biologically meaningful cell populations^37^. Together, these components allow the tissue to be modeled as a marked spatial point pattern, where points correspond to cell centroids and marks correspond to cell types^23^. This framework supports quantitative assessment of spatial organization beyond marginal cell-type abundances, enabling inference on whether specific cell populations exhibit non-random spatial structure.

To summarize spatial structure, established spatial summary statistics that characterize clustering, dispersion, and co-localization patterns are employed^25^. Single cell clustering summary measures, such as Ripley’s *K*-function and the nearest-neighbor distribution function *G*, are used to assess whether a cell type exhibits spatial clustering or dispersion relative to complete spatial randomness (CSR) at a given radius. Extensions of these statistics are used to evaluate spatial relationships between pairs of cell types, providing quantitative measures of attraction, repulsion, or spatial independence^25^.

The *spatialTIME* R package is a specialized framework for the visualization and quantitative analysis of mIF imaging data, designed to characterize both the abundance and spatial architecture of the TME^38,39^. The package represents mIF data as marked spatial point patterns, where individual cells are defined by their spatial coordinates and assigned phenotypes, and computes a range of spatial summary statistics, including Ripley’s *K*-function, nearest-neighbor based *G* statistics, and related measures of spatial association. These functions enable systematic assessment of cell-type specific clustering and cross-cell-type co-localization across tissue samples, with permutation-based inference to evaluate departures from spatial randomness.

FDA provides a framework for analyzing data observed as smooth functions over a continuous domain and has been widely used in fields such as longitudinal analysis, signal processing, and neuroimaging^40–42^. In FDA, an observed functional quantity *X_i_*(*r*), defined over a continuous domain *r* ∊ *R*, corresponding to spatial radius, is often represented using a basis expansion,

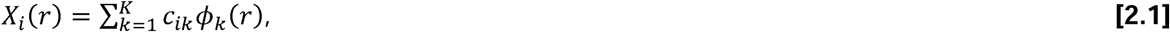

where *i* indexes subjects, 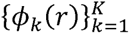 are basis functions, and *c_ik_* are subject-specific coefficients. In the context of spatial biology, FDA is particularly well suited for analyzing spatial summary statistics which are naturally evaluated over a continuum of spatial scales^37^. Treating these summaries as functional objects avoids reliance on a single, pre-specified radius and enables characterization of spatial behavior across the full range of distances^40,43^. For each subject, spatial clustering trajectories are calculated across a predefined range of radii for samples with enough positive cells of the phenotype of interest. When multiple tissue cores or ROIs are available per subject, subject-level spatial trajectories can be obtained by computing weighted averages of the estimated functions, with weights proportional to the number of cells within each tissue sample. Practical implementation of these steps, including smoothing, basis construction, and downstream modeling, can be performed using tools such as the R package *mxfda*^44^.

The *mxfda* R package is a specialized framework for the functional analysis of multiplex imaging data, designed to model spatial summary statistics as continuous functions and relate these spatial features to clinical outcomes. Building on spatial representations such as marked point patterns, *mxfda* facilitates the construction and smoothing of subject-specific spatial trajectories using flexible basis expansions and regularization techniques. The package provides tools for functional principal component analysis (FPCA), enabling decomposition of spatial trajectories into dominant modes of variation and extraction of low-dimensional functional features that summarize spatial organization across samples. In addition, *mxfda* supports regression modeling frameworks, including survival models, that incorporate functional predictors or their derived scores, allowing for joint assessment of spatial architecture and cell-type abundance. These capabilities enable a principled and scalable approach to quantify how multi-scale spatial patterns in the tumor immune microenvironment are associated with clinical outcomes.

FPCA was applied to the resulting spatial trajectories to obtain a low-dimensional representation of spatial variation across subjects. FPCA decomposes each spatial function *X_i_*(*r*) into a mean function *μ*(*r*) and a set of orthogonal eigenfunctions *ψ_k_*(*r*),

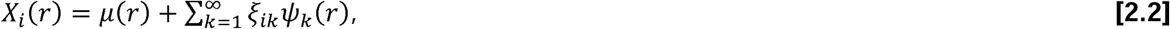

where *ξ_ik_* are the functional principal component (FPC) scores summarizing subject-specific deviations from the mean function^43,47^. These scores are defined as projections,

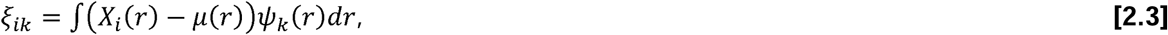

And capture dominant modes of variation across spatial scales. In this setting, the leading FPCs describe interpretable spatial patterns, such as overall clustering intensity or differential clustering across short- and long-range distances^45^. The estimated FPC scores were then used as covariates in downstream survival models, both as main effects and in interaction with cell-type abundance, enabling joint assessment of spatial architecture and cell abundance in relation to clinical outcomes.

### 2.3. Statistical Analysis

Spatial clustering trajectories were generated using the nearest-neighbor *G* function, as implemented in the *spatialTIME* R package. Analyses focused exclusively on *G*-based summaries, as we observed this spatial summary measure provided more interpretable associations with survival compared to Ripley’s *K*. These spatial trajectories were then analyzed using FDA, with functions centered relative to the CSR expectation to quantify departures from spatial randomness. Spatial summary functions were evaluated over a restricted range of radii from 5-100 pixels, corresponding to distances reliably assessed from TMA data. Spatial analyses were limited to the tumor compartment of TMAs or intratumoral ROIs. When both data on TMAs and ROIs were available for a given subject, ROIs were used (particularly for AACES).

Spatial trajectories were calculated only for tissue samples containing a minimum of six positive cells for each phenotype of interest, ensuring stable estimation of spatial summary measure. FPCA was applied to the resulting subject-level spatial trajectories, and the first two FPCs were retained, since they accounted for ∼84-97% of the variance for each study, for downstream modeling. The first FPC primarily captured overall clustering intensity across spatial scales, distinguishing tumors with globally higher versus lower levels of cell aggregation relative to CSR. The second FPC reflected scale-dependent variation in clustering, capturing differences in whether clustering occurred predominantly at shorter versus longer spatial distances. To facilitate comparison across studies, the direction of the FPCs was aligned so that the leading component corresponded to the same spatial pattern across studies. Representative mean-centered spatial trajectories and corresponding variation along the first two FPCs for CD8+ T cells across studies are shown in **Supplementary Figure S1**.

Associations between spatial clustering and risk of mortality were evaluated using Cox proportional hazards models. In the Cox proportional hazards models, the baseline hazard function *h*_0_(*t*) was left unspecified during model fitting, consistent with the partial likelihood formulation of the Cox model. Thus, inference for regression coefficients and hazard ratios (HRs) did not require explicit estimation of the baseline hazard. For adjusted survival curve visualization, study-specific baseline survival functions were subsequently obtained from the fitted Cox models using standard Breslow-type estimation.

For our primary analysis, we focus on the simplified Cox proportional hazards model defined in **Eq. 2.4**. A sequence of nested models was initially fit to evaluate the incremental contribution of spatial information, with preliminary models and corresponding meta-analytic results provided in the Supplementary Material. All models were adjusted for age at diagnosis, cancer stage [early vs. late], and cell-type abundance [low vs. high]. Abundance was dichotomized as high or low using a 1% threshold based on the proportion of positive cells within the tumor compartment, similar to prior ovarian cancer mIF study^27^. Cancer stage was categorized as “early” (stages I and II) or “late” (stages III and IV). Overall survival was defined as time from diagnosis to death from any cause, measured in months, with subjects censored at the date of last known follow-up if alive.

The final model focuses on *FPC*_1_ and its interaction with cell-type abundance. Based on the lack of association for *FPC*_2_ in the full interaction model, *FPC*_1_ was dichotomized to distinguish tumors with relatively low versus high spatial clustering, with values ≥ 0 classified as high clustering and values < 0 as low clustering. The resulting simplified interaction model is given by

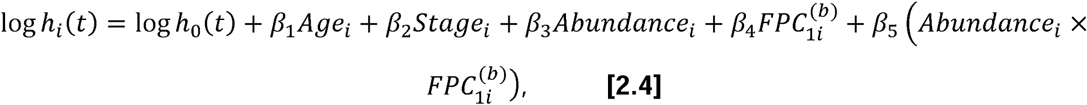

where 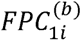 is 1 for low spatial clustering (low score) and 0 for high spatial clustering (high score). This model was used to estimate HRs and 95% confidence intervals for groups defined by combinations of abundance and spatial clustering, as well as to generate adjusted survival curves from the fitted Cox models.

All analyses were conducted separately within each study. Study-specific estimates were then combined using random-effects meta-analysis using the *meta*^46^ package in R to account for between-study heterogeneity, plotting results with forest plots. We used the *mxfda* R package to construct study-specific functional datasets of radius-indexed summary functions and perform FPCA to obtain per-sample FPC scores, which were then direction-aligned and used in the downstream Cox proportional hazards models. The proportional hazards assumption was assessed within each study using Schoenfeld residual-based diagnostics using the *survival*^47^ R package. Since this model incorporated spatial measures, subjects were restricted to those with sufficient cell counts for stable estimation of spatial trajectories, as shown in the last few rows of **Table 1**.

## 3. Results

In **Figure 1**, we present two representative samples from the NECC study illustrating contrasting spatial clustering patterns: low clustering (Sample A) and high clustering (Sample B). The top row shows the corresponding mIF images, highlighting overall tissue architecture and marker expression. The middle row displays the spatial distribution of cells by compartment, with tumor and stroma indicated and CD3+ T cells shown in red and CD8+ T cells in blue. In Sample A, T cells are sparsely distributed with minimal local aggregation, consistent with low abundance and a largely diffuse spatial pattern. In contrast, Sample B exhibits pronounced clustering, with distinct regions of densely aggregated T cells highlighted by green circles. These qualitative differences are quantitatively supported by the spatial clustering trajectories shown in the bottom row, which depict the degree of clustering across radii from 5-100 pixels. These visual and functional summaries illustrate the types of spatial patterns captured by the FDA-derived features used in downstream models. Values above zero indicate greater-than-random clustering, values below zero indicate spatial dispersion, and zero corresponds to CSR. Sample A shows values near zero across radii with a steady decrease as it gets closer to the 100-pixel radius, suggesting a pattern close to CSR with slight spatial dispersion. Sample B demonstrates consistently positive values, particularly from approximately 15 to 80 pixels, indicating strong clustering across multiple spatial scales. These findings highlight substantial heterogeneity in spatial organization across samples, motivating their integration into downstream survival modeling.

**Figure 1:**
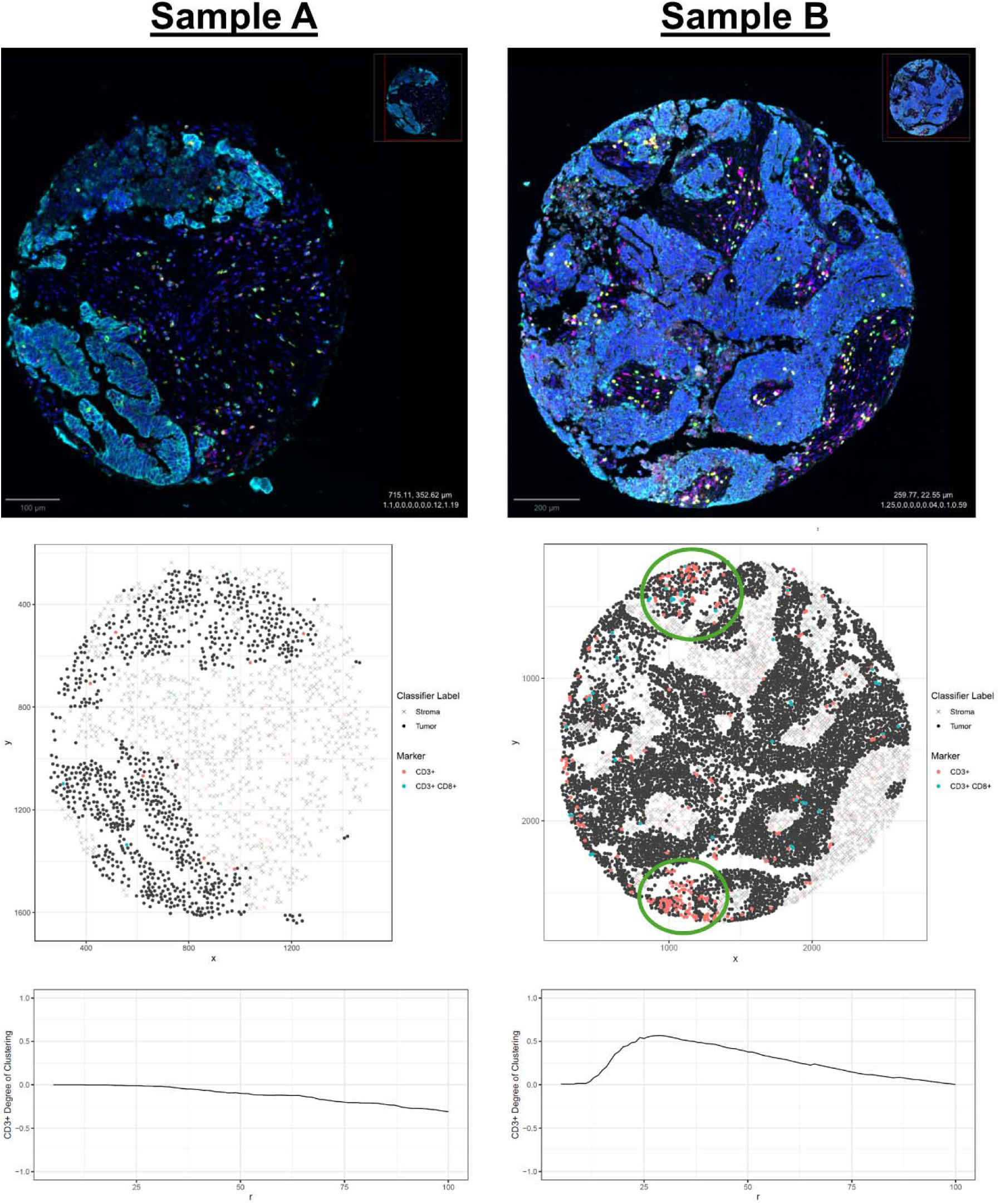
A low clustering sample (Sample A) and a high clustering sample (Sample B) from the NECC dataset with the raw mIF image (top row), plot of the tumor/stroma compartment cells with T cells and CD8+ T cells indicated (middle row), and the degree of clustering for those T cells across the 5-100 pixel-radius range (bottom row).

When assessing the proportional hazards assumption across studies, the estimated time-varying coefficients for the primary variables of interest (spatial clustering as captured by *FPC*_1_ and its interaction with abundance) were generally centered near zero with no consistent temporal trends, supporting the proportional hazards assumption. Some mild deviations were observed for certain covariates (e.g., stage and abundance) in individual studies, particularly at later follow-up times. However, these patterns were not consistent across studies and were accompanied by wide confidence bands, suggesting limited evidence of meaningful violation. A representative example is shown in **Supplementary Figure S2**, which displays the scaled Schoenfeld residuals for the *FPC*_1_ effect in AACES, illustrating the absence of a systematic time-dependent trend. Meta-analysis results for the preliminary models are shown in **Supplementary Figures S3-S7** and further discussed in Supplementary Materials.

**Figures 2** and **3** present the corresponding meta-analytic estimates for the primary model in this study (**Eq 2.4**). These forest plots display the study-specific and pooled estimates for the main and interaction effects, which form the basis for the group-specific HR estimates summarized in **Table 2**. Because interaction effects are most appropriately interpreted through joint consideration of both variables, we estimated HRs for groups defined by combinations of immune cell abundance (high vs. low) and spatial clustering (high vs. low), using low abundance and high clustering as the reference category (**Table 2**). For T cells, tumors characterized by high abundance and low spatial clustering exhibited lower estimated hazard relative to the reference group, whereas the group with high abundance and high clustering showed less favorable outcomes. Among CD8+ T cells, the survival advantage was most pronounced for tumors with high abundance and low spatial clustering (HR = 0.46, 95% CI: 0.34–0.61), while the group characterized by high abundance and high spatial clustering did not demonstrate comparable benefit. These findings indicate that the prognostic value of immune infiltration depends critically on its spatial organization, with more diffuse distributions associated with improved outcomes.

**Figure 2:**
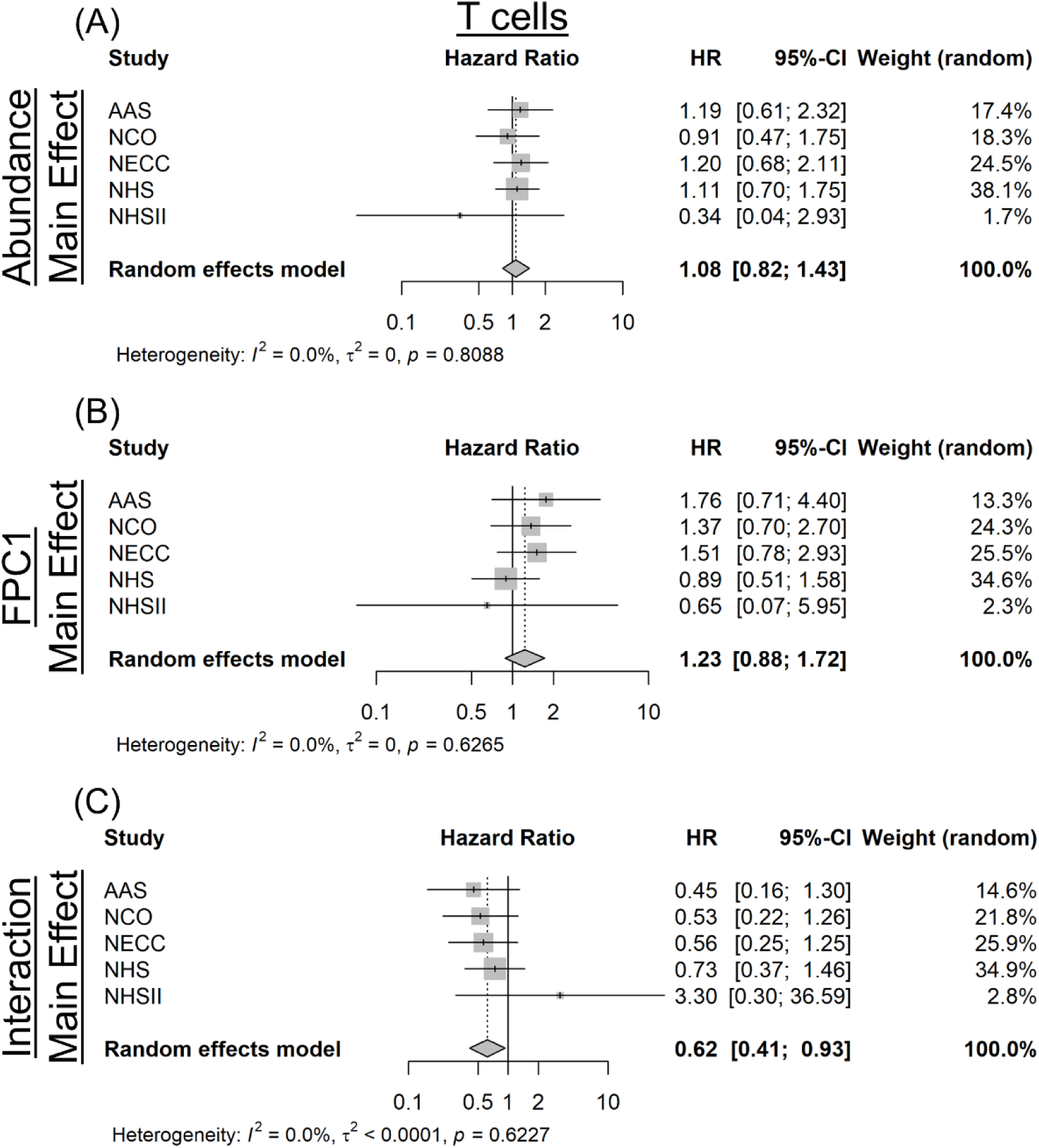
Forest plots showing study-specific and pooled HR estimates from the simplified interaction Cox model (**Eq. 2.4**) for T cells. Panels display (A) the main effect of abundance, (B) the main effect of spatial clustering (binary *FPC*_1_), and (C) the interaction between abundance and spatial clustering. Cell abundance was modeled as a binary variable (high vs. low, reference = low), and spatial clustering was defined as a binary variable based on the sign of the *FPC*_1_ score (high vs. low clustering, reference = high clustering). Models were adjusted for age at diagnosis and cancer stage (early vs. late, reference = late). Study-specific estimates were combined using random-effects meta-analysis.

**Figure 3:**
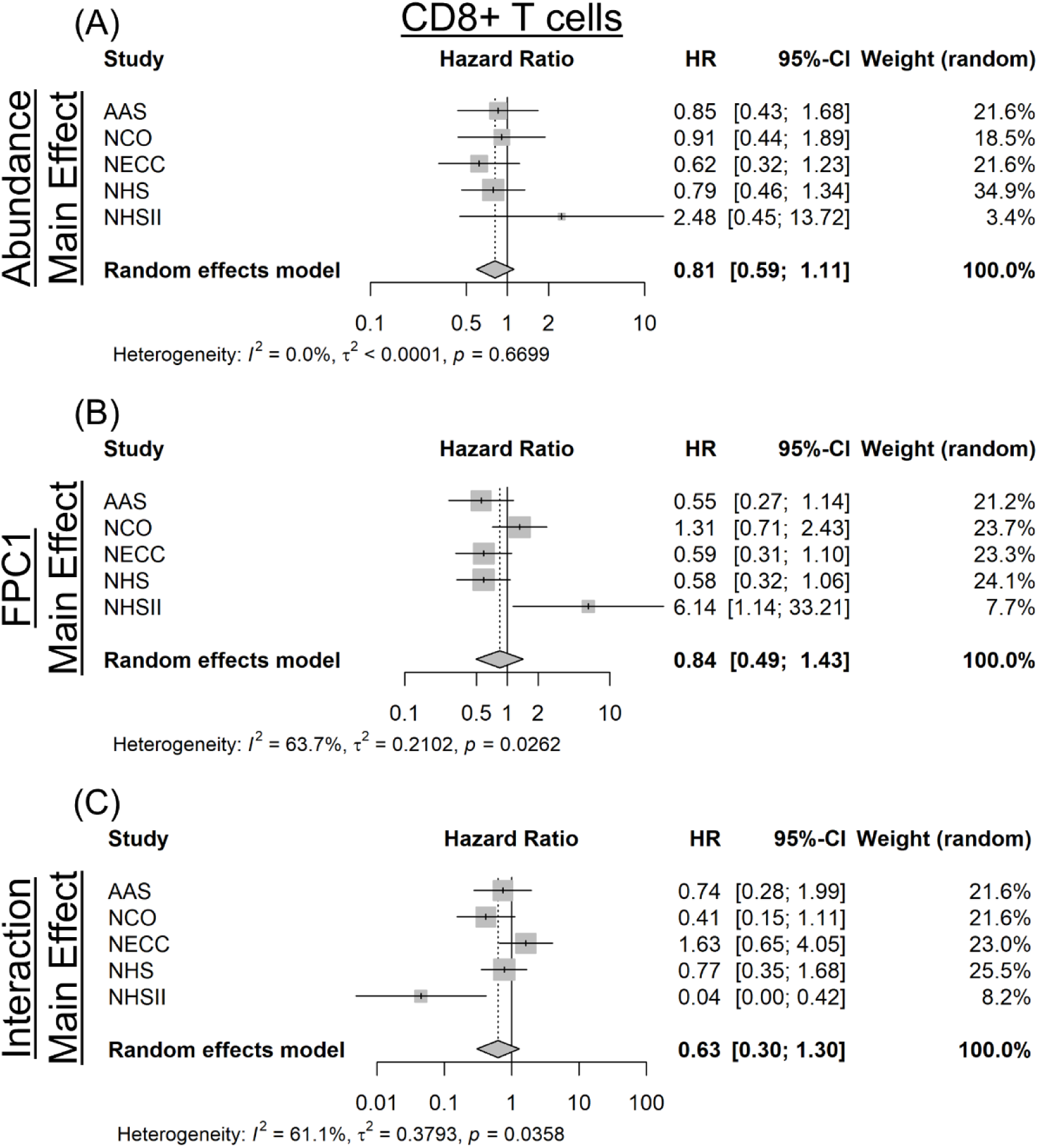
Forest plots showing study-specific and pooled HR estimates from the simplified interaction Cox model (**Eq. 2.4**) for CD8+ T cells. Panels display (A) the main effect of abundance, (B) the main effect of spatial clustering (binary *FPC*_1_), and (C) the interaction between abundance and spatial clustering. Cell abundance was modeled as a binary variable (high vs. low, reference = low), and spatial clustering was defined as a binary variable based on the sign of the *FPC*_1_ score (high vs. low clustering, reference = high clustering). Models were adjusted for age at diagnosis and cancer stage (early vs. late, reference = late). Study-specific estimates were combined using random-effects meta-analysis.

**Table 2:** HRs and 95% confidence intervals for each group defined by high or low abundance and spatial clustering (SC) for T cells (i.e., CD3+) and CD8+ T cells from the simplified interaction model (Eq 2.4).

| Cell Type | Group | HR | Lower CI | Upper CI | SE<sub>logHR</sub> |
| --- | --- | --- | --- | --- | --- |
| T cell | Low abundance + High SC | ref | ref | ref | ref |
|  | High abundance + High SC | 1.08 | 0.82 | 1.43 | 0.14 |
|  | High abundance + Low SC | 0.81 | 0.62 | 1.04 | 0.13 |
|  | Low abundance + Low SC | 1.23 | 0.88 | 1.72 | 0.17 |
| CD8+ T cell | Low abundance + High SC | ref | ref | ref | ref |
|  | High abundance + High SC | 0.81 | 0.59 | 1.11 | 0.16 |
|  | High abundance + Low SC | 0.46 | 0.34 | 0.61 | 0.15 |
|  | Low abundance + Low SC | 0.84 | 0.49 | 1.43 | 0.27 |

This pattern was further illustrated by Cox proportional hazard predicted survival curves displayed in **Figure 4**. The curves in **Figure 4** were generated from study-specific Cox models and pooled using the meta-analytic HR estimates, with age at diagnosis was fixed at the overall mean age at diagnosis across studies and stage set to high stage. These curves showed higher survival probabilities for only the high abundance, low clustering group for CD8+ T cells (**Figure 4B**). No abundance/clustering groups reached statistical significance for T cells, and corresponding survival curves exhibited this with little to no separation (**Figure 4A**).

**Figure 4:**
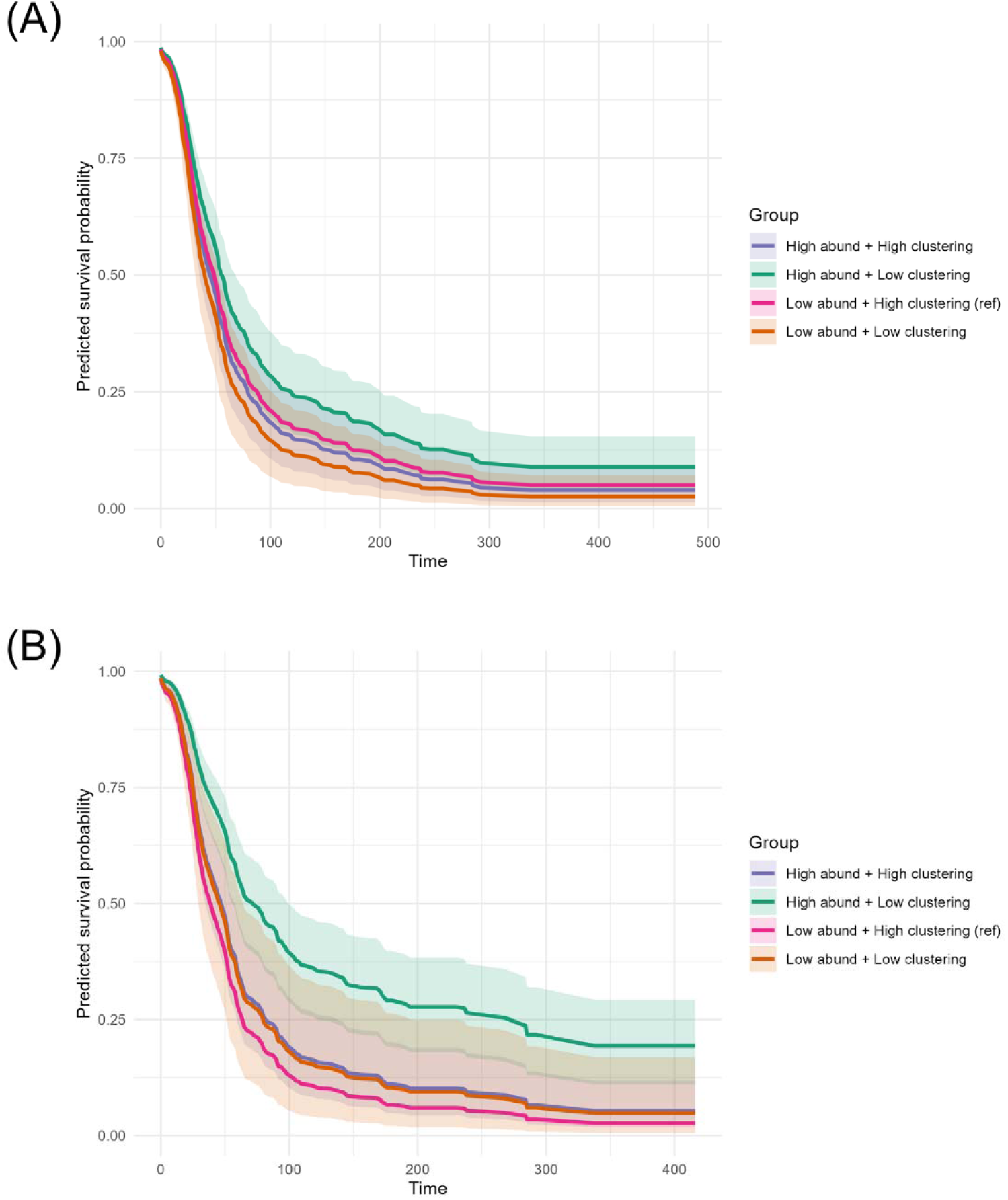
Adjusted model-based survival curves from the simplified Cox proportional hazards interaction model for (A) T cells and (B) CD8+ T cells.

## 4. Discussion

In this study, we proposed and applied an FDA framework to quantify and model spatial clustering of immune cells in the TME using mIF imaging data. By representing spatial summary statistics as trajectories across a continuum of spatial scales and summarizing them with FPCA, we avoided reliance on a single pre-specified radius and instead captured dominant modes of spatial organization. Applied across five ovarian cancer studies, this framework demonstrated that both immune cell abundance and spatial architecture contribute to overall survival, with evidence that spatial clustering modifies the prognostic effect of immune infiltration. These findings support the broader premise that spatial features of the ovarian TME provide clinically relevant information beyond cell abundance alone.

A key methodological contribution of this research is the flexibility of the FDA framework for spatial biology applications. Although the present analysis focused on within-cell-type clustering of T cells and CD8+ T cells using nearest-neighbor *G*-function trajectories, the same framework can be extended to other spatial summary functions that quantify cell-cell co-localization or multiscale spatial interactions. This adaptability makes the approach applicable to a wide range of biological questions involving tumor-immune interactions, stromal organization, or coordinated spatial patterning among multiple cell populations. In addition, because the framework operates on marked spatial point pattern representations, it is compatible with a variety of spatial profiling platforms, including imaging mass cytometry and spatial transcriptomics, enhancing its potential for broader application.

From a biological perspective, our findings are consistent with and extend prior work demonstrating that TILs are associated with improved outcomes in EOC^10–12^. Across studies, higher immune cell abundance was associated with improved survival, reinforcing the established role of CD8+ and broader lymphocytic infiltration in antitumor immunity. Importantly, our results indicate that abundance alone does not fully capture the prognostic relevance of the immune microenvironment. Spatial organization also plays a critical role, particularly for CD8+ T cells, where spatial features captured by the leading FPC were associated with survival. Across both T cell populations, increased spatial clustering was generally associated with poorer outcomes, with stronger and more precisely estimated effects observed for CD8+ T cells.

In the primary interaction analysis, tumors characterized by high immune cell abundance together with low spatial clustering exhibited the most favorable outcomes, particularly for CD8+ T cells. This suggests that a more spatially dispersed distribution of immune cells, rather than highly localized clustering, may reflect a more effective antitumor immune response. In the context of ovarian cancer, this pattern is biologically plausible, as a diffuse spatial distribution of CD8+ T cells may indicate broader tumor penetration and immune surveillance, whereas tight clustering may reflect localized immune confinement or spatially restricted immune activity. These results emphasize that the prognostic impact of immune infiltration depends on both the quantity and spatial organization of immune cells, and that these features should be interpreted jointly.

These findings are particularly relevant for ovarian cancer, where spatial context has long been hypothesized to influence prognosis but has often been summarized using fixed-radius statistics or coarse compartment-level measures. Prior studies have shown that the localization of T cells within tumor islets, rather than stromal regions, is associated with improved survival^10,11^, and potentially suggests that spatial immune features may help distinguish immunotherapy responders from non-responders. Our results extend this literature by demonstrating that the prognostic relevance of immune spatial organization is not limited to a single spatial scale. Instead, modeling clustering trajectories across radii provides a more comprehensive representation of multiscale spatial behavior and may better capture the biological complexity of the tumor immune microenvironment.

Several strengths of this study warrant emphasis. First, the multi-study design enhances generalizability and allows random-effects meta-analysis to account for between-study heterogeneity. Second, the FDA framework enables incorporation of multiscale spatial information without reducing the analysis to a single arbitrary distance, preserving richer structural detail. Third, the resulting low-dimensional representations remain interpretable and can be integrated into standard survival modeling frameworks.

In addition to these strengths, there are a few limitations. The analysis was restricted to subjects with sufficient spatial data and adequate cell counts for trajectory estimation (a requirement of utilizing FDA), leading to reduced sample size for spatially informed models. Additionally, because the data were derived from TMAs and intratumoral ROIs, the observed spatial patterns may not fully represent the broader tumor architecture. While nearest-neighbor *G*-function trajectories produced stable results in this setting, alternative spatial summary measures may capture complementary aspects of tissue organization and warrant further investigation. Notable heterogeneity in effect estimates was observed for the NHSII study, which differed from other studies in sample size and clinical characteristics, including a higher proportion of early-stage cases and younger age at diagnosis. These differences may lead to less stable estimates and highlight the importance of accounting for between-study variability in multi-study analyses.

Future work should explore extensions of this framework, including fully functional regression approaches that directly model spatial trajectories, as well as joint modeling of multiple cell types to capture coordinated spatial dynamics. From a translational perspective, further evaluation is needed to determine whether these spatial features improve prediction of response to chemotherapy or immunotherapy. Spatial features of CD8+ T cells may differ between immunotherapy responders and non-responders, suggesting that applying this framework in prospective clinical trial studies may be especially informative.

In summary, functional modeling of spatial clustering trajectories provides a flexible and interpretable approach for analyzing spatial biology data of the TME. By jointly accounting for immune cell abundance and spatial organization across multiple spatial scales, the proposed framework yields insights beyond traditional abundance-based or fixed-radius analyses and highlights the importance of spatial context in understanding tumor-immune interactions.

## Author Contributions

BLF conceived the project; JMS, ABL, LCP, SST, MKT, KLT generated the data; LCP, BMR quality controlled and organized the AACES & NCOCS data; KLT quality controlled and organized the NECC data; MKT quality controlled and organized NHSI/II data; BLF, LCP, ACS, JW developed the modeling framework; DL, ACS, BLF, CJS completed meta-analyses; BLF, CJS drafted the manuscript; all co-authors edited and reviewed the manuscript.

## Acknowledgments

We would like to thank the AACES investigators, Drs. Anthony Alberg, Elisa Bandera, Jill Barnholtz-Sloan, Melissa Bondy, Michele Cote, Ellen Funkhouser, Patricia Moorman, Edward Peters, Ann Schwartz, and Paul Terry, for their contributions to the AACES.

We would like to acknowledge the AACES interviewers, Christine Bard, LaTonda Briggs, Whitney Franx (North Carolina), Robin Gold (Detroit), and Myneka Macenat (New Jersey). We also thank the individuals responsible for facilitating case ascertainment across the 11 sites including Jennifer Burczyk-Brown (Alabama); Rana Bayakly and Vicki Bennett (Georgia); the Louisiana Tumor Registry; Lisa Paddock and Wendy Cedeno (New Jersey); Diana Slone, Yingli Wolinsky, Steven Waggoner, Anne Heugel, Nancy Fusco, Kelly Ferguson, Peter Rose, Deb Strater, Taryn Ferber, Donna White, Lynn Borzi, Eric Jenison, Nairmeen Haller, Debbie Thomas, Vivian von Gruenigen, Michele McCarroll, Joyce Neading, John Geisler, Stephanie Smiddy, David Cohn, Michele Vaughan, Luis Vaccarello, Elayna Freese, James Pavelka, Pam Plummer, William Nahhas, Ellen Cato, John Moroney, Mark Wysong, Tonia Combs, Marci Bowling, and Brandon Fletcher (Ohio); Martin Whiteside (Tennessee); and Georgina Armstrong and the Texas Registry, Cancer Epidemiology and Surveillance Branch, Department of State Health Services. We would like to thank Rex C. Bentley and Anne M. Mills for the review of the pathology in the AACES.

The authors would like to acknowledge the contribution to the NHS and NHSII from central cancer registries supported through the Centers for Disease Control and Prevention’s National Program of Cancer Registries (NPCR) and/or the National Cancer Institute’s Surveillance, Epidemiology, and End Results (SEER) Program. Central registries may also be supported by state agencies, universities, and cancer centers. Participating central cancer registries include the following: Alabama, Alaska, Arizona, Arkansas, California, Colorado, Connecticut, Delaware, Florida, Georgia, Hawaii, Idaho, Indiana, Iowa, Kentucky, Louisiana, Massachusetts, Maine, Maryland, Michigan, Mississippi, Montana, Nebraska, Nevada, New Hampshire, New Jersey, New Mexico, New York, North Carolina, North Dakota, Ohio, Oklahoma, Oregon, Pennsylvania, Puerto Rico, Rhode Island, Seattle SEER Registry, South Carolina, Tennessee, Texas, Utah, Virginia, West Virginia, Wyoming. The authors would also like to acknowledge the Channing Division of Network Medicine, Department of Medicine, Brigham and Women’s Hospital as home of the Nurses’ Health Studies. The authors assume full responsibility for analyses and interpretation of these data.

We would like to acknowledge the Advanced Analytical and Digital Laboratory under the Pathology Department at the Moffitt Cancer Center for the generation of the mIF data in the AACES and NHSI/II (Carlos Moran Segura, Jonathan Nguyen, Tony Alleyne, and Neale Lopez-Blanco).

## Code Availability

Code utilized in this research for the meta-analyses and survival analysis can be found on GitHub at https://github.com/FridleyLab/Ovarian_Cancer_FDA. This repository includes code using the following R packages: *meta*, for performing random-effects meta-analysis; *survival*, for fitting Cox proportional hazards models and conducting proportional hazards diagnostics; *spatialTIME*, for computing spatial summary statistics (e.g., nearest-neighbor *G*-functions) from multiplex imaging data; and *mxfda*, for constructing spatial trajectories, performing functional principal component analysis, and integrating functional features into downstream models.

## Data Availability

The AACES and NCOCS data is available for use by contacting J. Schildkraut at. The NECC data is available for use by contacting Kathryn Terry at. Because of participant confidentiality and privacy concerns, NHS/NHSII data are available upon reasonable written request. According to standard controlled access procedure, applications to use NHS/NHSII resources will be reviewed by an External Collaborators Committee for scientific aims, evaluation of the fit of the data for the proposed methodology, and verification that the proposed use meets the guidelines of the Ethics and Governance Framework and the consent that was provided by the participants. Investigators wishing to use NHS/NHSII data are asked to submit a description of the proposed project (https://www.nurseshealthstudy.org/researchers) or contact for details.

