## Supplemental Materials for "Functional Data Analysis of Spatial Clustering Identifies Prognostic T Cell Patterns in Ovarian Cancer"

This document contains the supplemental methods and results, including supplemental figures of results from the meta-analysis of the FPCA Cox models.

### 1. Statistical Models

All preliminary models were adjusted for age at diagnosis, cancer stage, and cell-type abundance, dichotomized as high or low using a 1% threshold based on the total number of cells in the tumor compartment of the tissue. The first model included all subjects (shown in the top row of **Table 1**) and served as a baseline model without spatial trajectories. This model is exhibited as

$$\log h_i(t) = \log h_0(t) + \beta_1 Age_i + \beta_2 Stage_i + \beta_3 Abundance_i + \beta_4 S.b_i, \quad \textbf{[S1.1]}$$

where $Age_i$ is the age at diagnosis of subject $i$, $Stage_i$ is the binary cancer stage variable of subject $i$, $Abundance_i$ is the dichotomized cell-type abundance variable of subject $i$ with a 1% threshold, and $S.b_i$ is the binary indicator variable denoting whether spatial clustering could be estimated for subject $i$ (i.e., whether the minimum cell count criterion of 6 cells of a specific population was met). This formulation allowed inclusion of all tumors while accounting for potential differences between tumors with and without spatial data.

The second model was restricted to subjects with 6 or more positive cells to compute spatial trajectories and is written as

$$\log h_i(t) = \log h_0(t) + \beta_1 Age_i + \beta_2 Stage_i + \beta_3 Abundance_i + \beta_4 FPC_{1i} + \beta_5 FPC_{2i}, \quad \textbf{[S1.2]}$$

where $FPC_{1i}$ is the first FPC score for subject $i$, and $FPC_{2i}$ is the second FPC score for subject $i$. This model assessed whether spatial clustering patterns were associated with survival independently of cell abundance.

The third model extended the previous model by including interaction terms between cell-type abundance and each FPC score, exhibited as

$$\log h_i(t) = \log h_0(t) + \beta_1 Age_i + \beta_2 Stage_i + \beta_3 Abundance_i + \beta_4 FPC_{1i} + \beta_5 FPC_{2i} + \beta_6 (Abundance_i \times FPC_{1i}) + \beta_7 (Abundance_i \times FPC_{2i}), \quad \textbf{[S1.3]}$$

allowing the association between spatial clustering and survival to vary according to abundance. Inference from this model focused on contrasts comparing combinations of abundance and spatial clustering levels.

### 2. Results

We first evaluated the abundance-only model (**Eq. S1.1**) as a baseline framework that excluded spatial information, allowing assessment of the association between dichotomized immune cell abundance and overall survival while adjusting for age at diagnosis, cancer stage, and an indicator for the availability of spatial data. Across studies, higher abundance of both T cells (**Figure S3A**) and CD8+ T cells (**Figure S3B**) was associated with improved survival (T cells: HR = 0.81, 95% CI: 0.66–0.98; CD8+ T cells: HR = 0.64, 95% CI: 0.52–0.79). These effects were consistent in direction across most studies. However, the NHSII study showed notable deviations and contributed to between-study heterogeneity. This likely reflects differences in study characteristics, including a smaller sample size, a higher proportion of early-stage cases, and younger age at diagnosis.

We next examined whether spatial clustering patterns were associated with survival independently of cell abundance by introducing the first two FPC scores derived from nearest-neighbor *G*-function trajectories (model expressed in **Eq. S1.2**). Since this model and the subsequent models include spatial measures, analyses were restricted to subjects with sufficient spatial data as indicated in **Table 1**. For T cells, higher $FPC_1$ scores, corresponding to increased spatial clustering, were associated with higher estimated hazard (HR = 1.06, 95% CI: 0.99–1.14; **Figure S4A**), although the confidence interval included the null. Notably, the direction of effect was consistent with that observed for CD8+ T cells, for which higher $FPC_1$ scores were associated with significantly worse survival (HR = 1.17, 95% CI: 1.04–1.33; **Figure S4B**). These results suggest that increased spatial clustering is generally associated with poorer outcomes across T cell populations, with stronger effects observed for CD8+ T cells. No significant associations were observed for $FPC_2$ for either cell type (**Figure S5**), suggesting that the primary axis of spatial variation is the dominant contributor to prognostic signal.

To assess whether the effect of spatial clustering depended on immune cell abundance, we extended the model to include interaction terms between abundance and each FPC score (**Eq. S1.3**). **Figures S6** and **S7** summarize the main and interaction effects. For T cells, evidence of interaction between abundance and $FPC_1$ suggests that the association between spatial clustering and survival differs by abundance level. Rather than interpreting the interaction coefficient directly, the combined effects indicate that the prognostic impact of spatial clustering is modified by overall cell abundance. In particular, the estimated effects imply differing survival patterns across combinations of abundance and clustering, motivating further evaluation through group-based comparisons. For CD8+ T cells, higher abundance remained associated with improved survival, while higher $FPC_1$ scores were associated with worse survival. Together, these findings suggest that the relationship between immune infiltration and clinical outcome depends jointly on both the quantity and spatial organization of immune cells.

### Supplemental Figure Legends

**Figure S1:** Mean-centered *G*-function trajectories (black) and deviations along the first two FPC ($FPC_1$ and $FPC_2$) are shown across radii of 5-100 pixels for each study (rows: AACES, NCOCS, NECC, NHS, NHSII). Blue and red curves indicate positive and negative deviations along each FPC, respectively.

**Figure S2:** Assessment of the proportional hazard assumption for $FPC_1$ in AACES (T cells). Scaled Schoenfeld residuals are shown with a smoothed estimate of the time-varying coefficient and corresponding confidence bands.

**Figure S3:** Forest plots showing study-specific and pooled HR estimates for the effect of cell-type abundance on overall survival for (A) T cells and (B) CD8+ T cells, based on the abundance-only Cox proportional hazards model (**Eq. S1.1**). Cell abundance was modeled as a binary variable (high vs. low), defined using a 1% threshold based on the proportion of positive cells within the tumor compartment, with low abundance serving as the reference group. Models were adjusted for age at diagnosis, cancer stage (early vs. late, reference = late), and an indicator for the availability of spatial data. Study-specific estimates were combined using random-effects meta-analysis.

**Figure S4:** Forest plots showing study-specific and pooled HR estimates for the main effect of $FPC_1$ from the Cox model including spatial features (**Eq. S1.2**) for (A) T cells and (B) CD8+ T cells. $FPC_1$ represents the first FPC score derived from nearest-neighbor *G*-function trajectories, treated as a continuous variable. Models were adjusted for age at diagnosis, cancer stage (early vs. late, reference = late), and cell-type abundance (high vs. low, reference = low). Study-specific estimates were combined using random-effects meta-analysis.

**Figure S5:** Forest plots showing study-specific and pooled HR estimates for the main effect of $FPC_2$ from the Cox model including spatial features (**Eq. S1.2**) for (A) T cells and (B) CD8+ T cells. $FPC_2$ represents the first FPC score derived from nearest-neighbor *G*-function trajectories, treated as a continuous variable. Models were adjusted for age at diagnosis, cancer stage (early vs. late, reference = late), and cell-type abundance (high vs. low, reference = low). Study-specific estimates were combined using random-effects meta-analysis.

**Figure S6:** Forest plots showing study-specific and pooled HR estimates from the Cox model with interaction terms (**Eq. S1.3**) for T cells. Panels display (A) the main effect of abundance, (B) the main effect of $FPC_1$, and (C) the interaction between abundance and $FPC_1$. Cell abundance was modeled as a binary variable (high vs. low, reference = low), and $FPC_1$ was treated as a continuous variable. Models were adjusted for age at diagnosis and cancer stage (early vs. late, reference = late). Study-specific estimates were combined using random-effects meta-analysis.

**Figure S7:** Forest plots showing study-specific and pooled HR estimates from the Cox model with interaction terms (**Eq. S1.3**) for CD8+ T cells. Panels display (A) the main effect of abundance, (B) the main effect of $FPC_1$, and (C) the interaction between abundance and $FPC_1$. Cell abundance was modeled as a binary variable (high vs. low, reference = low), and $FPC_1$ was treated as a continuous variable. Models were adjusted for age at diagnosis and cancer stage (early vs. late, reference = late). Study-specific estimates were combined using random-effects meta-analysis.

**Supplemental Figure 1:**

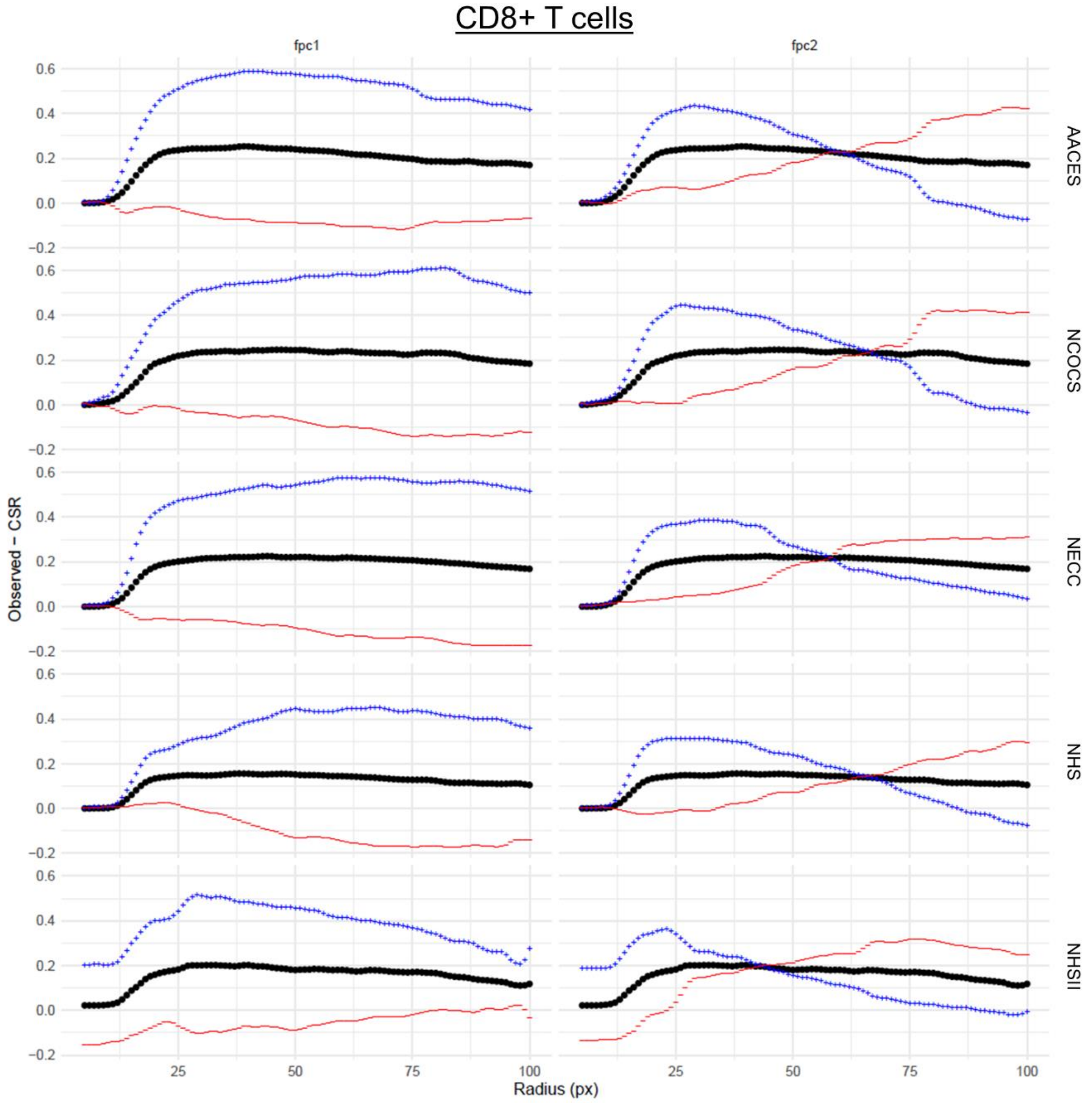

**Supplemental Figure 2:**

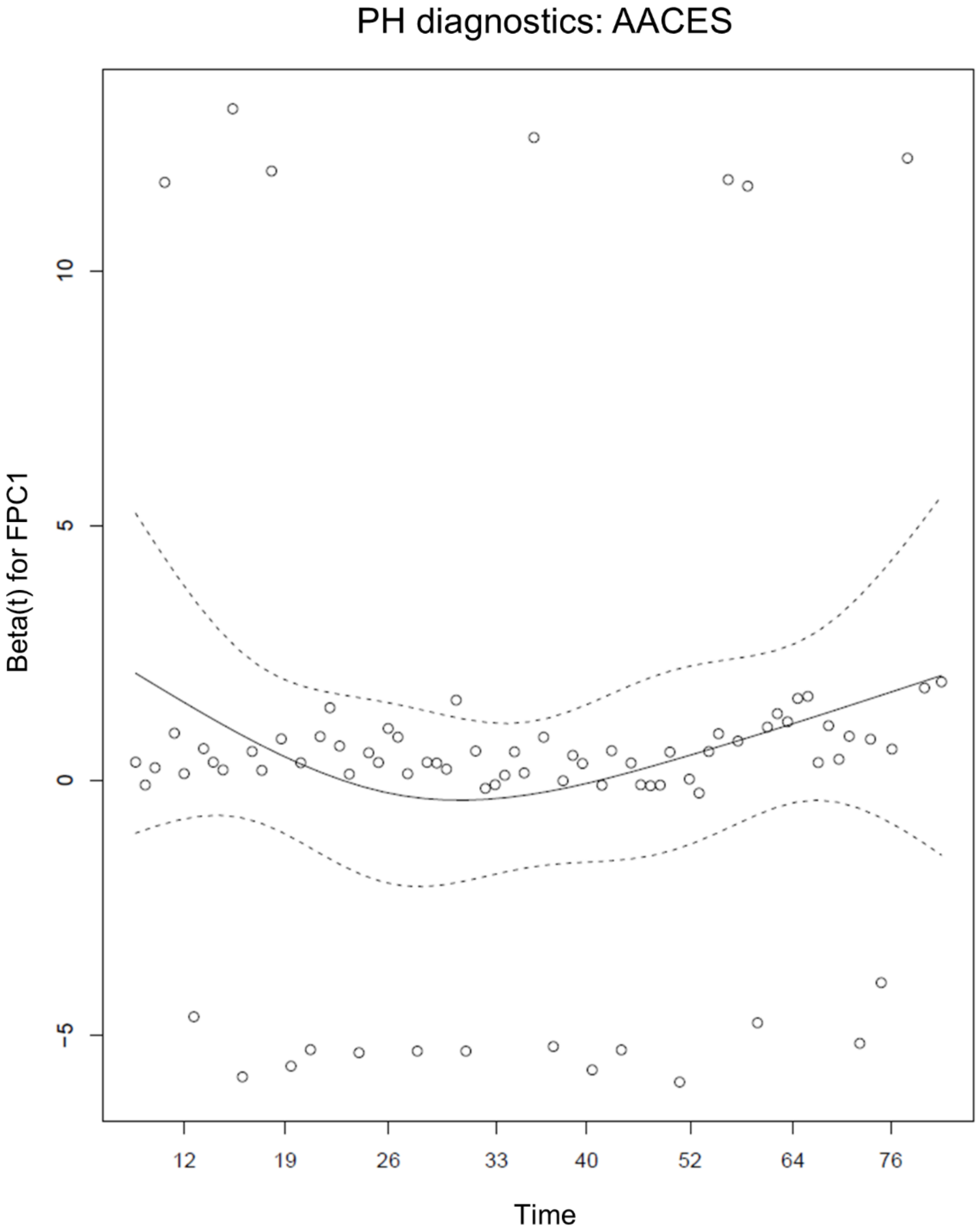

**Supplemental Figure 3:**

(A) T cells

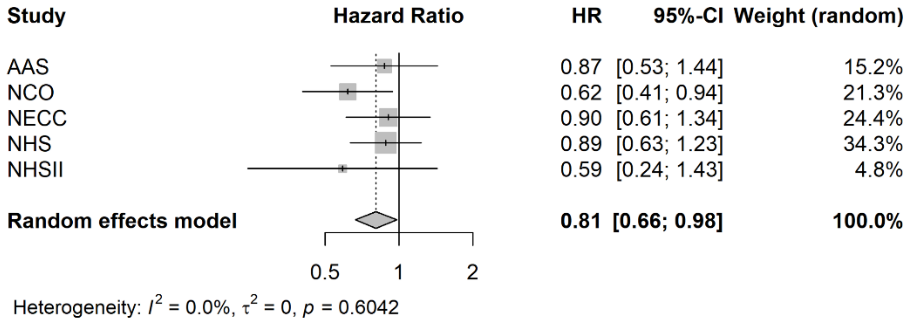


(B) CD8+ T cells

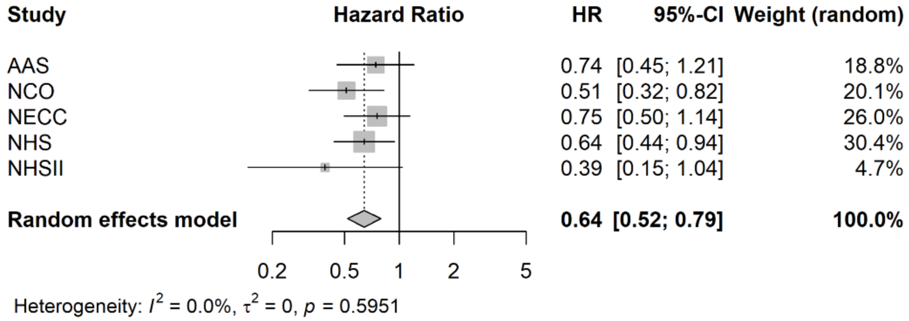

**Supplemental Figure 4:**

(A) T cells

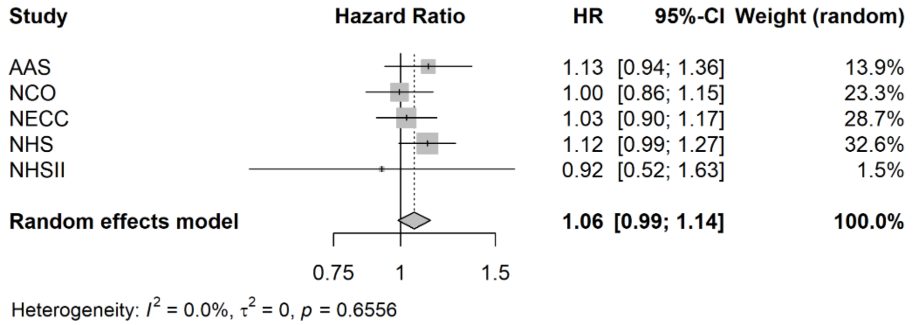


(B) CD8+ T cells

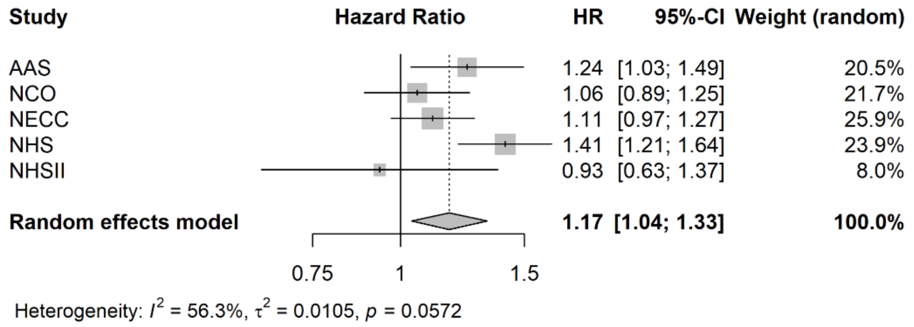

**Supplemental Figure 5:**

(A) T cells

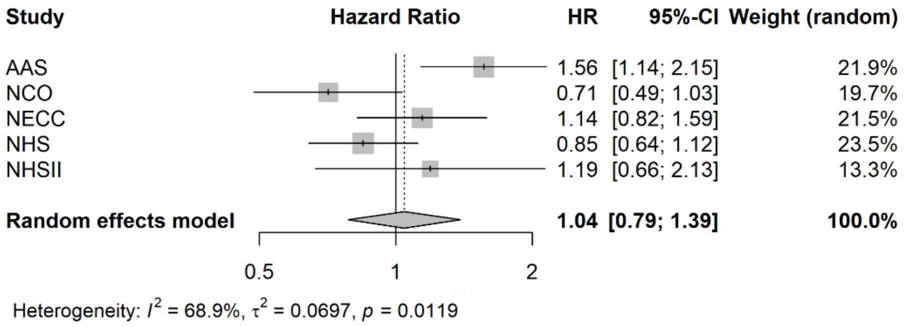


(B) CD8+ T cells

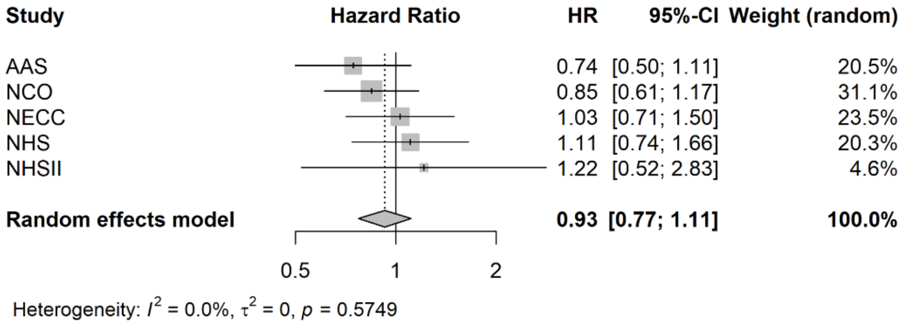

**Supplemental Figure 6:**

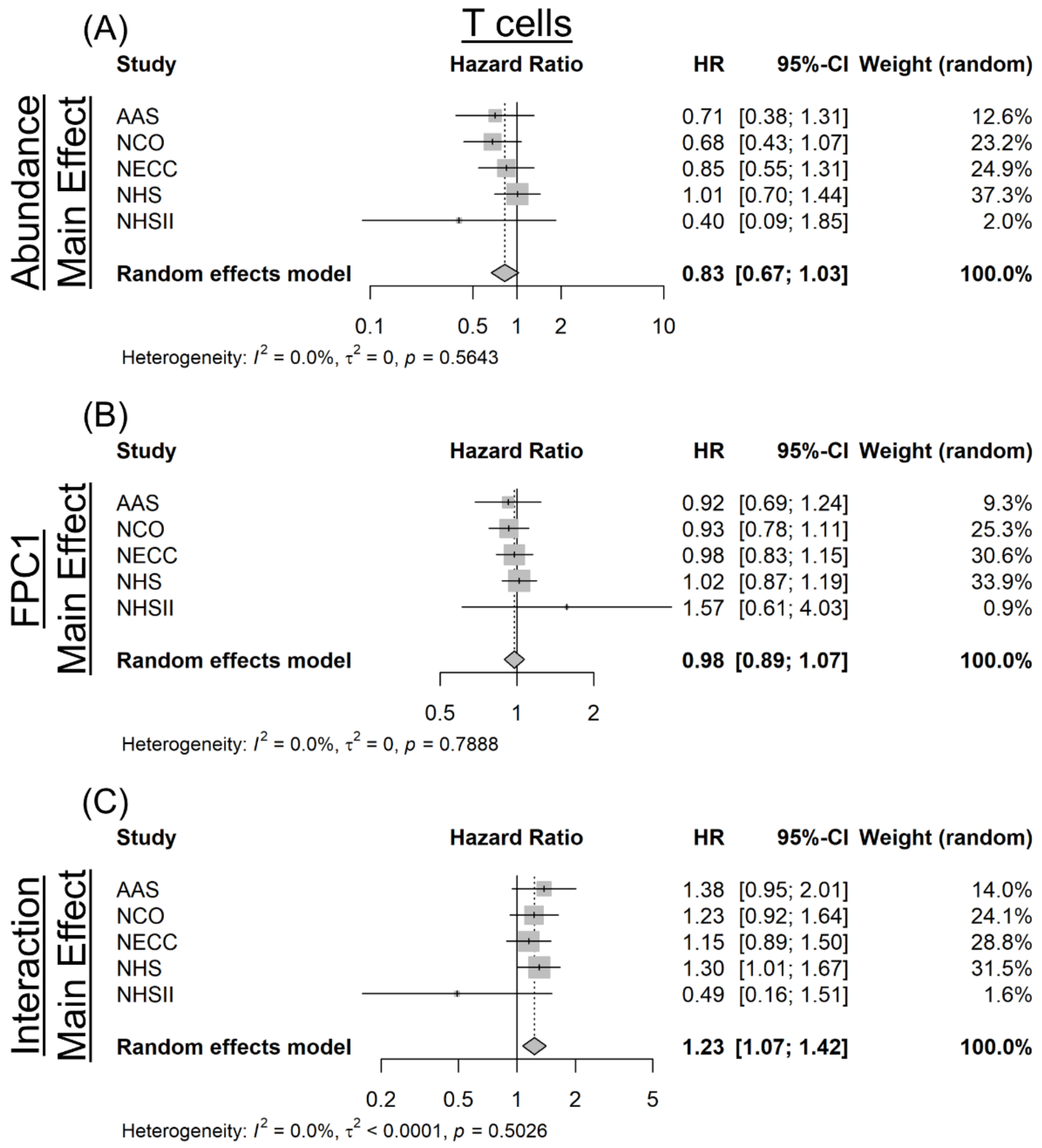

**Supplemental Figure 7:**

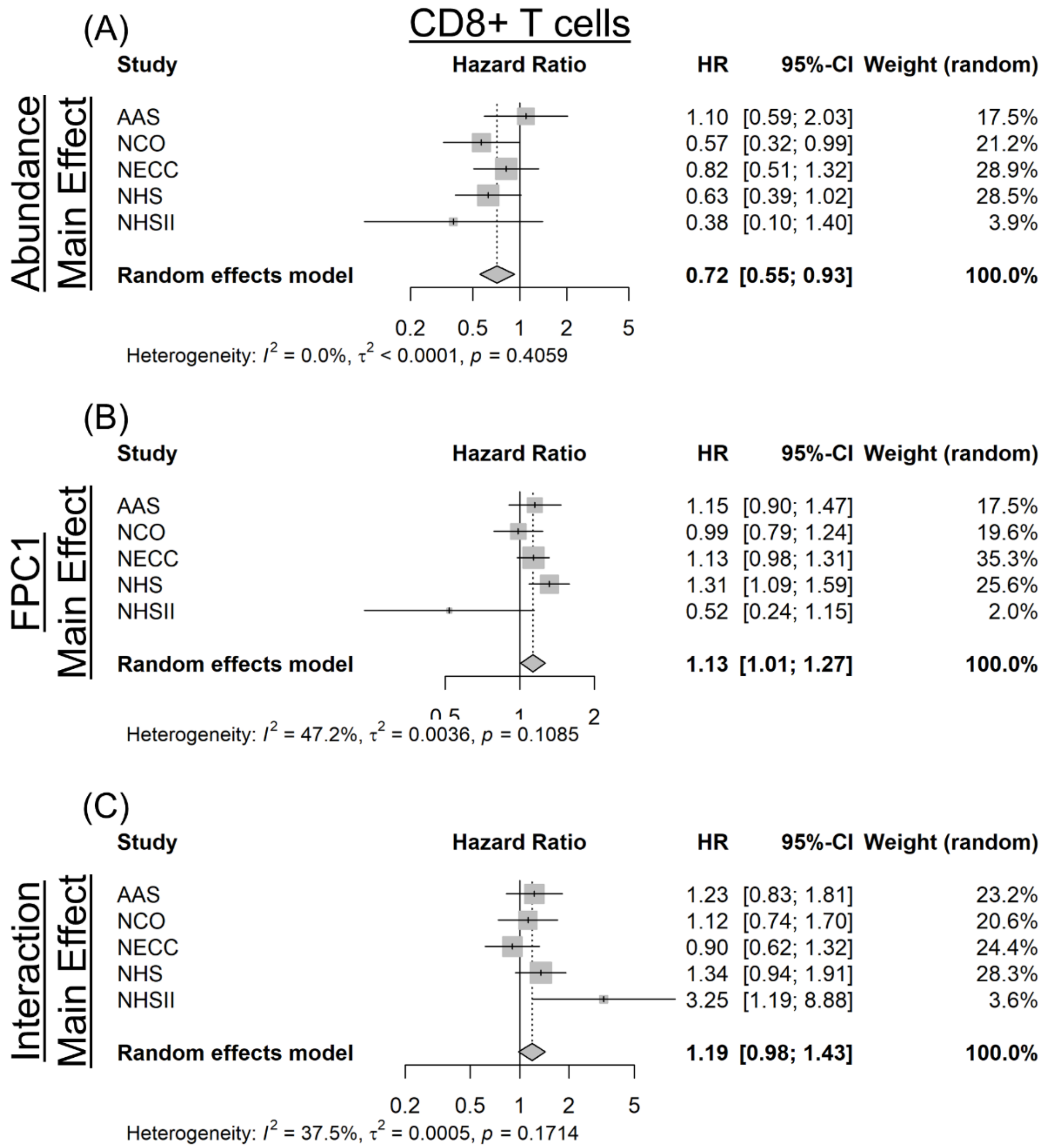
